# Reversible inhibition of viral life cycle in response to elevated temperature in a bloom-forming alga

**DOI:** 10.64898/2026.06.02.729564

**Authors:** Noa Shima, Talia S. Shaler, Nir Joffe, Daniella Schatz, Assaf Vardi

## Abstract

Ocean warming is expected to reshape microbial interactions and community composition, with profound consequences for marine biogeochemical cycles. Host–virus dynamics are central to these processes, yet their response to elevated temperature remains poorly understood. Here, using the bloom-forming alga *Gephyrocapsa huxleyi* and its specific large dsDNA virus, *Emiliania huxleyi* virus (EhV), we show that elevated temperature withheld virion production and abolished distinct stages of the viral life cycle. Viral adsorption and early transcription remained active, whereas viral DNA replication and late gene expression were arrested, leading to an intracellular inhibition of infection. Intriguingly, this arrested infection state was reversible following prolonged heatwave conditions. Single-cell analyses revealed that reversibility of infection inhibition occurred both within a small fraction of infected cells and by reinfection by extracellular virions that remained viable during heat exposure in the extracellular milieu. Furthermore, we detected variability in infection inhibition by temperature across several host-virus pairs, suggesting that the effect of temperature is strain-specific. Our findings uncover a temperature-sensitive checkpoint in the viral life cycle, providing a mechanistic framework for understanding how marine heatwaves may reshape virus-driven mortality and carbon cycling in the ocean.

## Introduction

The ocean uptakes approximately 90% of Earth’s excess heat caused by anthropogenic greenhouse gas emissions [1]. It is estimated that over the next century, sea surface temperatures will increase by 1–6°C and exhibit greater fluctuations due to climate change [2, 3]. Accompanied by the overall increase in water temperature, events of extreme heat, described as marine heatwaves, have already increased and will continue to rise in frequency, intensity, and duration [4]. While many studies explore the effects of individual or combined stressors on specific species or community composition, a major knowledge gap remains on how climate change will alter microbial interactions, including host-virus interactions in the marine environment.

Phytoplankton are unicellular, photosynthetic microorganisms that form the basis of marine food webs and contribute about half of the estimated global net primary production [5, 6]. Lytic viruses are central to controlling algal bloom termination worldwide, diverting carbon flux from higher trophic levels to bacterial respiration and deep ocean sequestration [7, 8]. Blooms of the cosmopolitan coccolithophore *Gephyrocapsa huxleyi* are often terminated following infection by the lytic large double-stranded DNA coccolithovirus, the *Emiliania huxleyi* virus (EhV) [9, 10]. During infection, EhV reprograms *G. huxleyi* to support production of new virions, producing a unique metabolic state of the infected host cell, termed the “virocell” [11, 12]. The EhV infection cycle has been proposed to comprise five temporal transcriptional classes, from the immediate-early to the late kinetic class [13]. Successful infection requires viral attachment and entry to the algal cell, transcription, translation, DNA replication, formation of progeny virus particles, and release from the host cell. Replication of EhV is proposed to occur in the nucleus, while assembly and genome packaging are thought to take place in the cytoplasm, with mature virions acquiring an outer membrane and exiting the cell [14]. Disrupting any of these steps may lead to an ineffective or unsuccessful infection, enabling the host cell to survive.

There is substantial intraspecific diversity among *G. huxleyi* strains in their thermal response mechanisms, with thermal niche width ranging from 6°C to 25°C, highlighting the capacity of this diverse group to survive across a range of temperatures [15, 16]. Nevertheless, Elevated temperatures can alter phytoplankton bloom dynamics and affect the cellular proteome of key phytoplankton species, with temperatures beyond their thermal optima inducing heat stress response and reducing the efficiency of carbon fixation, potentially reducing their contribution to the oceanic carbon sink [3, 16-19]. An increase in temperature is likely to affect viruses as well, as their life cycle is closely linked to host metabolism [20]. For example, the virulence of *Heterosigma akashiwo* virus is temperature-dependent, displaying unsuccessful infection below or above the optimal temperature range of 20 to 25°C [21]. Decay rates of natural virus populations in the western Pacific Ocean have been shown to increase at a +4°C temperature shift [22]. In *Micromonas* virus, infection shifts from a classical lytic mode to a chronic-like life cycle at elevated temperatures [23]. Changes in the susceptibility of *G. huxleyi* to its virus (EhV) at warmer temperatures have already been observed and attributed to changes in the host cell surface lipid composition that can block EhV adsorption, although no precise mechanism has been identified [24].

Here, we show that the infection process of EhV is suppressed intracellularly at an elevated temperature. We demonstrate that the virus successfully adsorbs to and enters host cells but cannot replicate its genome or complete gene transcription. Infection arrest lasts for an ecologically relevant time scale, the duration of a marine heatwave, and is reversible upon return to basal temperature. Lastly, we demonstrate that the infection outcome under heat varies across different EhV-*G. huxleyi* pairs, suggesting that future ocean warming will shape viral-host interactions in the marine environment.

## Materials and Methods

### Culture collection and maintenance

*G. huxleyi* RCC6918, RCC6936, RCC6946, and RCC6962 strains were isolated from a mesocosm experiment performed by our lab in Bergen, Norway, 2018 [25]. *G. huxleyi* CCMP2090, CCMP374 and CCMP379 strains were obtained from the National Center for Marine Algae. All cultures were maintained in filtered seawater (FSW) supplemented with f/2 (-Si) (Guillard and Ryther, 1962) at 18°C with a 16:8 h light cycle, under a light intensity of 100 μmol photons m^2^ s^-1^ provided by cool white LED lights. EhV-M1 (hereon EhVM1) was isolated from the mesocosm experiment by propagation on *G. huxleyi* CCMP374. For the experiments in this manuscript, EhVM1 was propagated on *G. huxleyi* RCC6946 by infecting an exponentially growing culture and filtering the lysate through a 0.2 µm polycarbonate filter. Viruses were concentrated using an Amicon Ultra Centrifugal Filter, 100 kDa MWCO. All experiments were performed at an ambient temperature of 18°C or an elevated temperature of 23°C, with a 16:8 h light cycle, under a light intensity of 70 µmol photons m^2^ s^-1^ provided by cool white LED lights. The freshly prepared EhVM1 stock was inoculated into *G. huxleyi* cultures at the early exponential growth phase with a multiplicity of particles (MOP) of 3 and supplemented with Ampicillin or Kanamycin (100 μg ml^-1^ or 50 μg ml^-1^, respectively)

### Quantification of algal abundance, viral abundance, and cell death

Flow cytometry analyses were conducted using a CytoFLEX S Flow Cytometer (Beckman Coulter) equipped with four lasers: blue (488 nm), red (638 nm), violet (405 nm), and yellow (561 nm). *G. huxleyi* abundance: quantified using flow cytometry or plate reader. Flow cytometry-*G. huxleyi* cells were identified by plotting chlorophyll fluorescence (excitation at 561 nm, emission at 665–715 nm) against side scatter (as a proxy for cell size) and quantified by counting the high chlorophyll events. Plate reader-the algal population density was evaluated using an Infinite 200 PRO multimode plate reader (Tecan Group Ltd.), by measuring algal chlorophyl autofluorescence (excitation at 480 nm, emission at 680 nm). Virus enumeration: samples were fixed with 0.5% glutaraldehyde for at least 30 minutes at 4°C, plunged into liquid nitrogen, and stained with SYBR gold (Invitrogen) diluted 1:10,000 in Tris-EDTA buffer. Then, samples were incubated for 20 minutes at 80°C and cooled to room temperature. Samples were then analyzed by flow cytometry (excitation at 488 nm, emission at 500–550 nm). Algal cell death measurement: samples were stained with 1 μM of the nucleic acid stain SYTOX Green (Invitrogen), which only enters cells with compromised membranes. The stained samples were incubated in the dark at room temperature for 20 minutes and analyzed by flow cytometry (excitation at 488 nm, emission at 500–550 nm). An unstained sample was used as a control for the background signal.

### Growth rate calculation

Growth rates (μ_max_) were calculated during exponential growth of uninfected cultures at each temperature, using the natural logarithm of cell abundance: μ = (ln N_2_ − ln N_1_) / (t_2_ − t_1_), where N represents cell concentration at time t.

### Quantification of photosynthetic efficiency

Photosynthetic efficiency (Fv/Fm) was measured using a Water-PAM fluorometer (WALTZ) [26]. Fm represents the maximum fluorescence emission level in the dark measured with a saturating pulse of light (450 nm, 2700 μmol photons m^−2^ s^−1^, 800 ms), Fv = Fm–F0.

### Assessing virus adsorption to *G. huxleyi* cells

To assess viral adsorption to *G. huxleyi* RCC6946, cells were grown at 18°C to the late exponential phase (∼3 × 10^6^ cells ml^-1^), concentrated by 5000g x 5 min centrifugation, and divided into six biological replicates. Half of the samples (n=3) were placed at 23°C 15 min prior to infection, while the other half remained at 18°C. Following this incubation, all replicates were infected with EhVM1 at an MOP of 0.1. Immediately after infection, each of the three biological replicates at both temperatures was divided into 21 1.5 ml tubes (three technical repeats per time point), resulting in 63 tubes per temperature, each containing 500 µl of infected culture. Extracellular (unattached) viruses were counted at 2, 15, 30, 60, 90, 120, and 180 minutes post-infection. At each time point, samples were centrifuged at 3000g X 3 min, and the supernatant containing unattached viruses was transferred to a new 1.5 ml tube. Viruses were fixed in glutaraldehyde and counted as described above. The percentage of adsorbed viruses equals 100% minus the percentage of unattached viruses.

### RNA harvesting and reverse transcription quantitative PCR

To quantify viral transcription, RT-qPCR was performed on infected cultures, using primers targeting viral genes from different kinetic classes (e.g., Early and Late genes) (Table 1). At each time point, 50 ml of algal culture was taken from each biological replicate, centrifuged at 3000 x g for 10 min at 4°C, the supernatant was discarded, and the remaining volume was centrifuged again at 5000 x g for 5 min at 4°C, followed by removal of the remaining supernatant. The pellet was then resuspended in 450 μl of RLT buffer supplemented with 1% β-Mercaptoethanol, plunged into liquid nitrogen and stored at -80°C until RNA extraction. To disrupt the cells, glass beads (≤106 µm, Sigma-Aldrich) were added to the pellets, and the samples were placed in a bead beater and run at 30 Hz for 5 min. RNA was extracted using the RNeasy Plant Mini Kit according to the manufacturer’s instructions. To remove genomic DNA contamination, RNA was treated with Turbo DNase (Ambion) in four rounds of enzyme addition and incubation at 37°C. DNase was removed using RNAClean XP beads. RNA concentration was measured using Qubit. cDNA (3 ng µl^-1^) was prepared using Verso cDNA Kit. For the no-RT controls, cDNA reactions (1.5 ng µl^-1^) were prepared without reverse transcriptase for each sample.

**Table 1.**
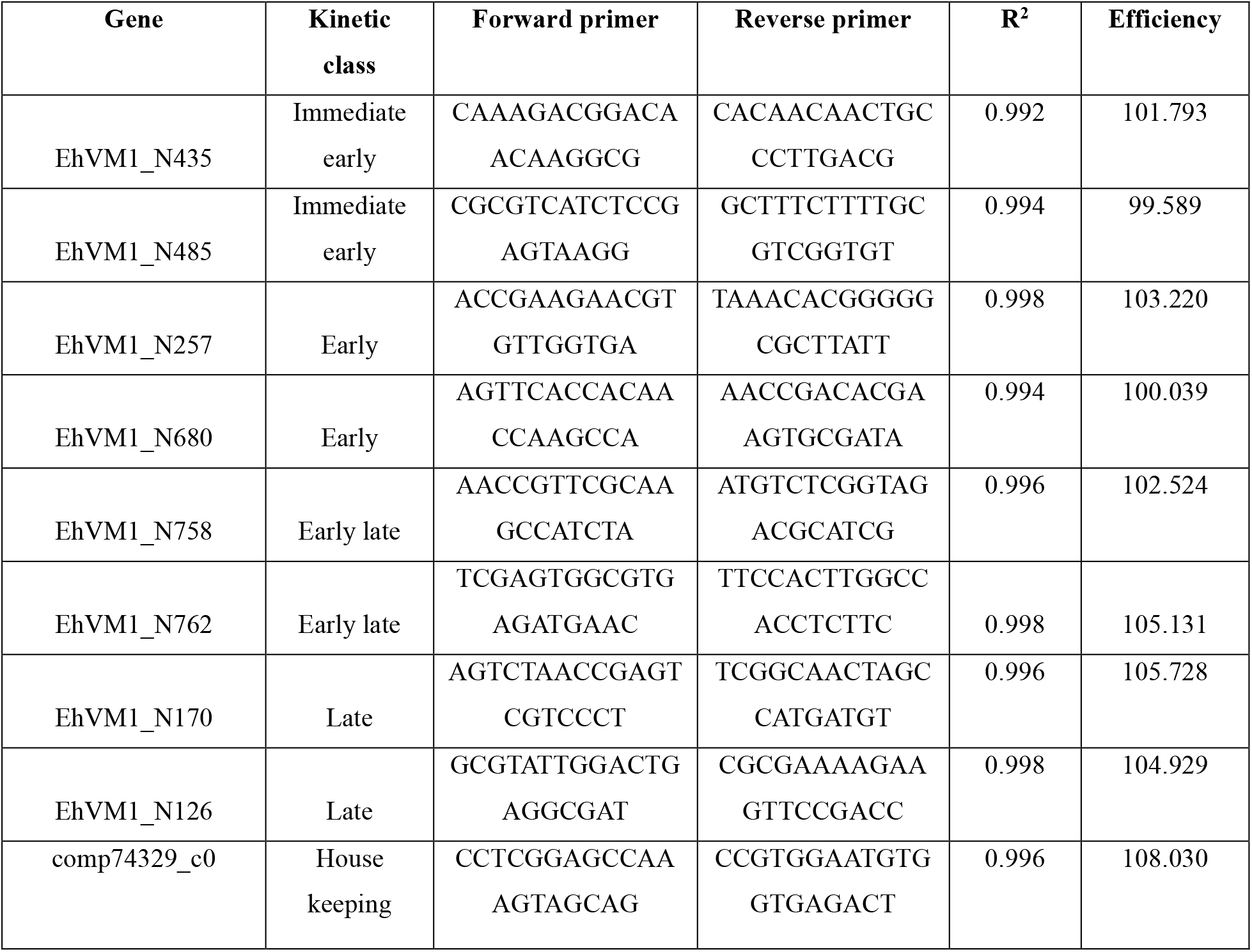
Primers for RT-qPCR.

To assign predicted kinetic classes to EhVM1 genes, we used the kinetic class annotation of EhV201 genes (genome: JF974311) established by Ku et al.,2020 [13], based on their temporal expression (immediate-early, early, early-late, and late). The reference EhVM1 genome has 489 predicted coding sequences [27], to which we added 306 potential ORFs (795 transcripts in total). Genes from the same kinetic class were shown to share specific promoter motifs, which are conserved across multiple EhV strains [13]. For the purposes of the present analysis, all late classes (late 1, late 2 and late-undefined) were combined into a single “late” category because they all represent genes expressed during the late stages of infection and the same promoter motif. Putative EhVM1 orthologs of the EhV201 genes were identified using BLASTP. For each EhVM1 gene, a 200 bp sequence window centered on the translation start codon (±100 base pairs from the first base of ATG) was extracted following the approach of Ku et al., 2020 [13]. The four kinetic class-specific promoter motifs described by Ku et al. were searched against these sequences using FIMO (Find Individual Motif Occurrences) from MEME Suite v5.5.0 (fimo --thresh 1.0E-4, Table S1) [28]. Genes were considered representative markers of a given kinetic class when (i) their closest EhV201 homolog belonged to that kinetic class and (ii) the corresponding kinetic class-specific promoter motif was detected. Candidate marker genes were further filtered using bulk RNA-seq data from *G. huxleyi* RCC6946 infected with EhVM1 previously obtained in our lab [29]. Only genes with sufficient expression levels (baseMean > 40) were retained. For late genes, we additionally required significant differential expression at 24 h or 48 h post-infection. A housekeeping gene was selected from the same RNA-seq dataset as a gene showing stable expression across all infection time points (baseMean > 150; −0.5 < log_2_FC < 0.5). Additionally, the chosen gene showed no differential expression in another bulk RNA-seq comparing exponentially growing and stationary-phase *G. huxleyi* RCC6946 [29].

### Digital droplet PCR

Digital droplet PCR was used to quantify viral DNA replication. At each time point, 3 ml of each sample was centrifuged at 3000 x g for 3 min. The supernatant was discarded, and pellets were washed three times with filtered sea water (FSW). Then, the pellets were plunged into liquid nitrogen and stored at -80°C. DNA was extracted using REDExtract-N-Amp plant PCR kit (Sigma-Aldrich). EhV-mcp-FAM and Ghux-18S-HEX probes were designed using BioRad’s program (Table 2). PCR reactions (20 µl) contained 10 µl 2× Supermix for probes, 1 µl of each probe, 1 µl HaeII restriction enzyme, 1 µl template DNA diluted 20-fold, and nuclease-free water. PCR was performed with an initial enzyme activation at 95°C for 10 min, followed by 40 cycles of denaturation at 94°C for 30 sec, annealing at 57°C for 1 min, and enzyme deactivation at 98°C for 10 min. All reactions were performed on the QX200 Droplet Digital PCR system.

**Table 2.**
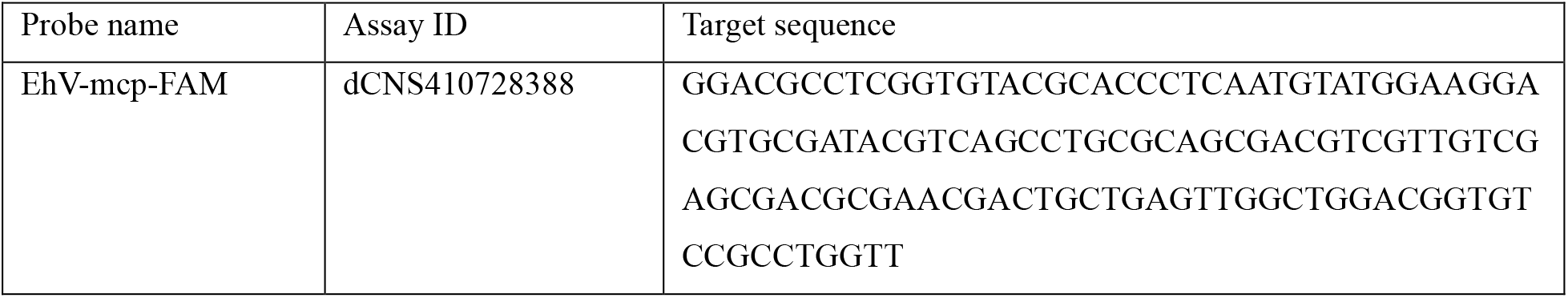
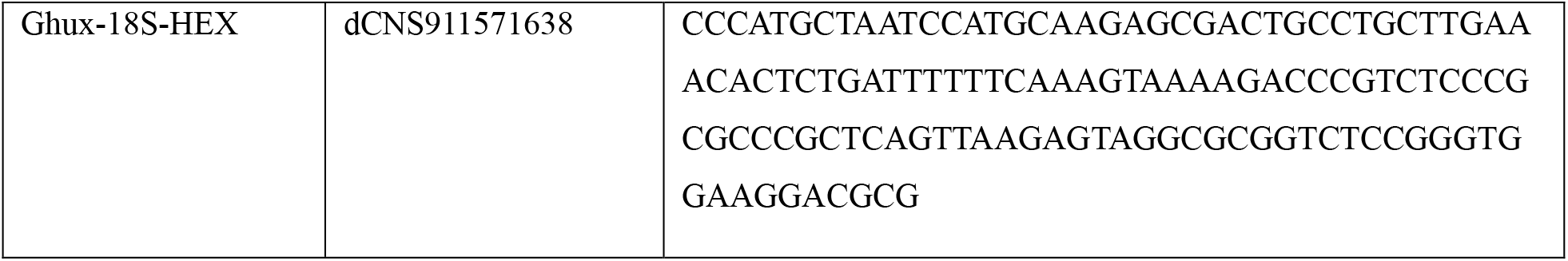
Probes for digital droplet PCR.

### Testing viral infectivity derived from single virocells

To test reactivation of infection after heat exposure at the single-cell level, we applied a single-cell infectivity assay modified from Kirsner et al., 2016 [30]. *G. huxleyi* RCC6946 was infected with EhVM1 and incubated at 18°C or 23°C for 22 hours. Extracellular viruses were removed by washing algal samples twice by centrifugation (3000 × g, 5 min), discarding the supernatant, and resuspending the pellets in 30 ml FSW. Subsequently, single cells were sorted using a BD FACS Aria-III into 96-well plates containing 200 µl of exponentially growing RCC6946, with the addition of Ampicillin and Kanamycin (100 μg ml^-1^ and 50 μg ml^-1^, respectively). Single cells were sorted into each well based on their optical properties (chlorophyll, forward light scattering, and high green autofluorescence, Fig. S3A). The plates were wrapped with parafilm to prevent evaporation and placed at 18°C for 6 days under low light (∼20 µmol photons m^-2^ s^-1^). Wells were considered lysed if the fluorescence signal (measured using the plate reader, as described above) was < 500 arbitrary units (AU, average of control non-lysed wells was >5000 AU). Lysis of cells in acceptor wells indicated that the sorted cells were virocells that completed the infection cycle upon return to 18°C.

### Statistical analysis

Statistical differences in growth rates were calculated using one-way ANOVA. Statistical differences in DNA replication were calculated using Welch’s t-test. Statistical differences for the single-cell sorting experiment were calculated using logistic regression, with well lysis (fluorescence signal <= 500 AU) treated as a binary variable (lysed=0, alive=1). All other statistical differences were evaluated using two-way repeated measures (ANOVA), followed by Tukey’s or Sidak’s post-hoc test. A mixed-design model (split-plot ANOVA) was used whenever data were missing. GraphPad Prism 9.3.1. was used to generate graphs and perform statistical analysis.

## Results

### Viral infection is suppressed at elevated temperatures

We first explored how elevated temperature affects viral infection. The *G. huxleyi* strain RCC6946 was infected with EhVM1 at a multiplicity of infection (MOP) of 3 (three viruses per cell) and placed at either 18°C, 21°C, 23°C, or 25°C for 7 days. Uninfected cultures were also placed at each temperature as controls. The maximal algal growth rate of control cultures at 18°C was 0.74 day^-1^ and was not significantly affected by increasing temperatures (Fig. S1). At 18°C, *G. huxleyi* RCC6946 was susceptible to EhVM1. Within 7 days post-infection (hereafter, dpi), algal abundance and photosynthetic yield declined by 9,200-fold (p<0.0001) and 3-fold (p=0.0266) compared to the 18°C uninfected control, respectively (Fig. 1A-B). The percentage of dead cells in the population, assessed by Sytox Green, increased to 76% at 7 dpi, 10-fold higher than in the control (p<0.0001, Fig. 1C). Viral abundance increased 232-fold compared to 0 dpi (p=0.0337, Fig. 1D). An increase in temperature gradually reduced viral infection. At 21°C, at 7 dpi, algal abundance, photosynthetic efficiency, and percentage of cell death of infected cultures were not statistically different from those at 18°C, but viral production was reduced 12-fold (p=0.0295, Fig.1). Algal abundance in the infected cultures at 23°C and 25°C was three orders of magnitude higher than at 18°C at 7 dpi (p=0.005 for 23°C, and p=0.0133 for 25°C, Fig. 1A). Photosynthetic efficiency was 3-fold higher than 18°C infection (p=0.0248 for 23°C and p=0.0226 for 25°C) (Fig. 1B). Compared to 18°C, algal cell death was 8-fold lower at 23°C and 5-fold lower at 25°C (p<0.0001, Fig.1C). Notably, at no time point was the viral concentration at 23°C or 25°C statistically different from that at 0 dpi (Fig. 1D). As 23°C was the lowest temperature at which the infection was completely arrested, we chose it as a model temperature to investigate the mechanisms governing infection suppression at elevated temperatures.

**Figure 1.**
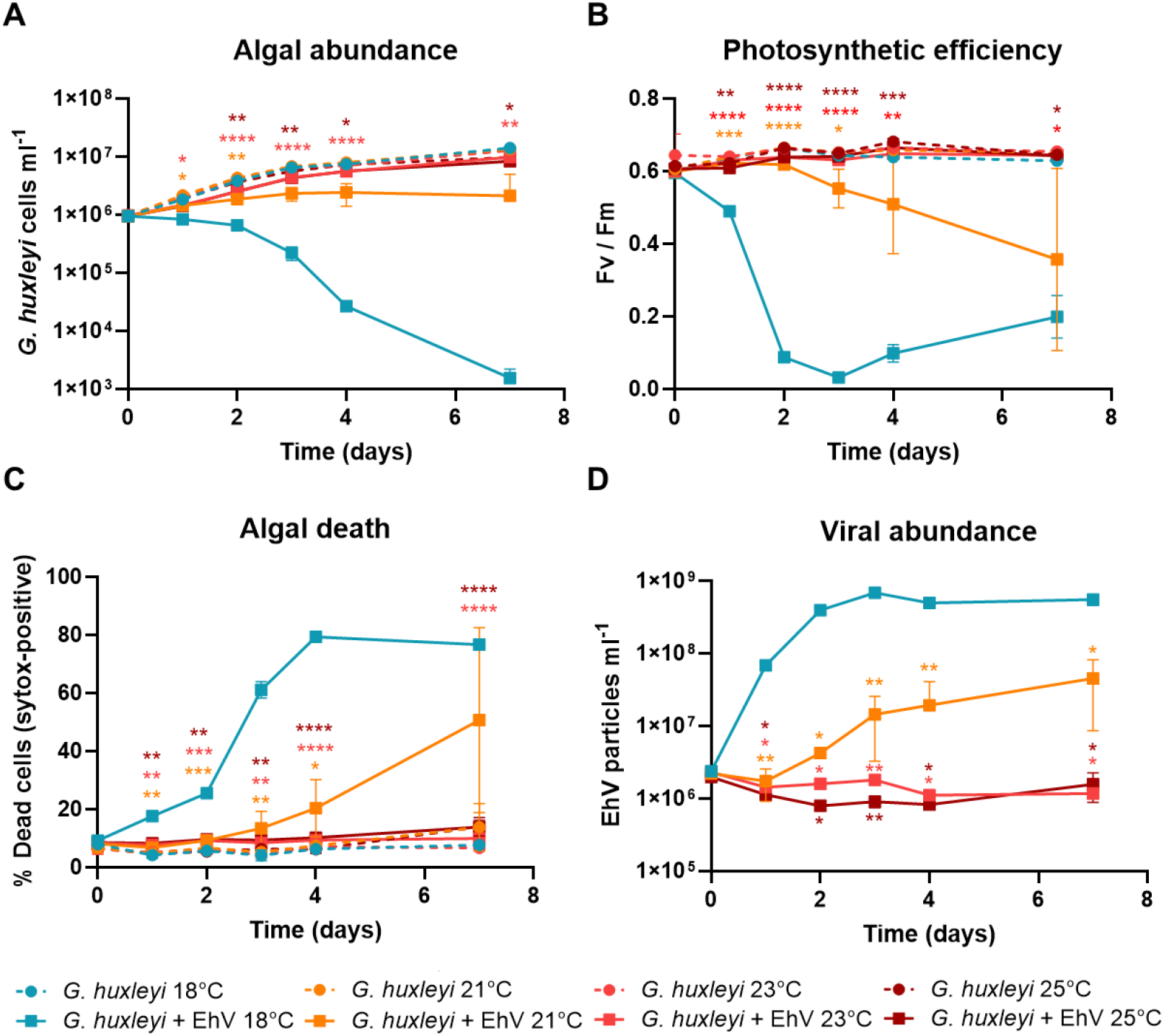
Viral infection is suppressed at elevated temperatures. *G. huxleyi* strain RCC6946 was infected with EhVM1 (solid lines, squares) and placed at either 18°C (blue), 21°C (orange), 23°C (red), or 25°C (dark red) for 7 days. Uninfected cultures were also placed at each temperature as controls (dashed lines, circles). **A)** Algal cell abundance. **B)** Photosynthetic efficiency as assessed by measuring the maximum quantum yield of photosystem II (Fv/Fm), calculated as (Fm − F0)/Fm, using pulse-amplitude modulated (PAM) fluorometry. **C)** Algal cell death as quantified by the percentage of cells positively stained with the cell death marker SYTOX-Green. **D)** Extracellular EhV abundance in infected cultures at all temperatures. Results for A, C, and D were evaluated by flow cytometry and are presented as the average ± SD (n = 3). Error bars smaller than symbol size are not shown. Statistical differences were tested using repeated-measures two-way ANOVA followed by Tukey’s multiple comparisons test. Colored asterisks indicate the statistical difference between the infected culture at each temperature and the 18°C-infected culture. p < 0.05 (*), p < 0.01 (**), p < 0.001 (***), p < 0.0001 (****).

### Elevated temperature affects infection on the cellular level

To differentiate between the impact of temperature on the virion infectivity versus the host cell, we pre-incubated the algal host and the virus separately at 23°C for 3 days before infection at the standard temperature of 18°C (Fig. 2A). As a positive control for successful infection, *G. huxleyi* RCC6946 was inoculated with EhVM1 at 18°C without pre-incubation at 23°C of either the virus or the algae. Inoculation of the 23°C-treated virions with 23°C-treated algae did not abolish infection at 18°C. There was a slight delay in algal demise compared to the untreated cultures at 71 (p=0.0009) and 77 (p=0.0380) hours post-infection (hereafter, hpi), but by 102 hpi, the algal abundance in the treated culture was reduced to similar levels as the control infection (Fig. 2B). In accordance, viral abundance increased without a significant difference compared to the control infection (Fig. 2C). As viral infection was successful with both heat-treated viruses and heat-treated algal cells, we concluded that viral infectivity and host susceptibility are not abolished at 23°C. We thus hypothesized that sensitivity to elevated temperature occurs during the life cycle of EhV.

**Figure 2.**
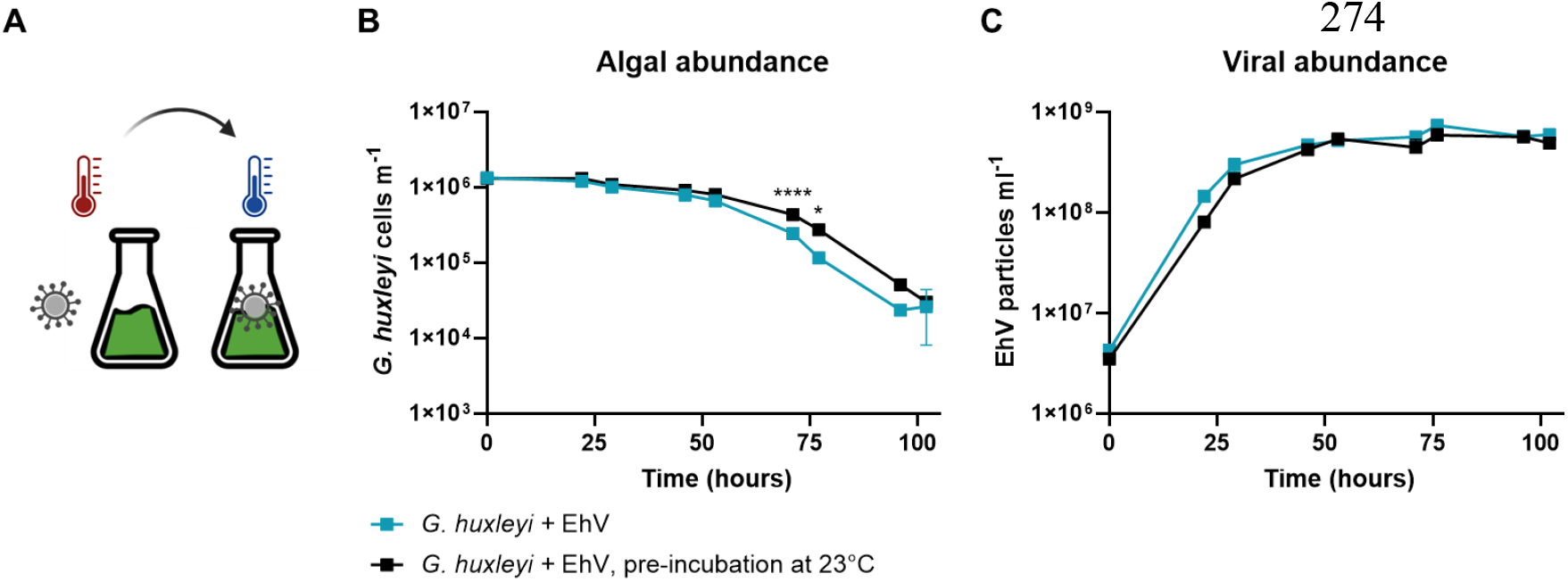
Heat pretreatment does not abolish viral infectivity or algal sensitivity to infection. **A)** Schematic representation of the experimental system. EhVM1 and *G. huxleyi* RCC6946 were incubated separately at 23°C for 72 hours prior to infection at 18°C (black squares). Non-heat-treated viruses and algal cells served as the control for 18°C infection (blue squares). Algal abundance **(B)** and viral abundance **(C)** were evaluated by flow cytometry and represent average ± SD (n = 3). Error bars smaller than symbol size are not shown. Statistical differences were tested using repeated-measures two-way ANOVA followed by Sidak’s multiple comparisons test. p < 0.05 (*), p < 0.0001 (****). The scheme presented in (A) was created with Biorender.com.

### Viral attachment is successful under elevated temperature

We aimed to pinpoint the exact phase in the infection cycle that is hindered at an elevated temperature (Fig. 3A). We first assessed whether EhV attachment to cells is compromised at 23°C, which could account for the observed infection arrest. We performed an adsorption assay to compare the attachment of EhVM1 to the surface of *G. huxleyi* RCC6946 cells at both temperatures. Surprisingly, elevated temperature did not modulate viral adsorption, and the dynamics were similar at both temperatures, with no significant difference in the fraction of viruses attached to cells. At 90 minutes post-infection, ∼80% of viruses were adsorbed to the cells (Fig. 3B).

**Figure 3.**
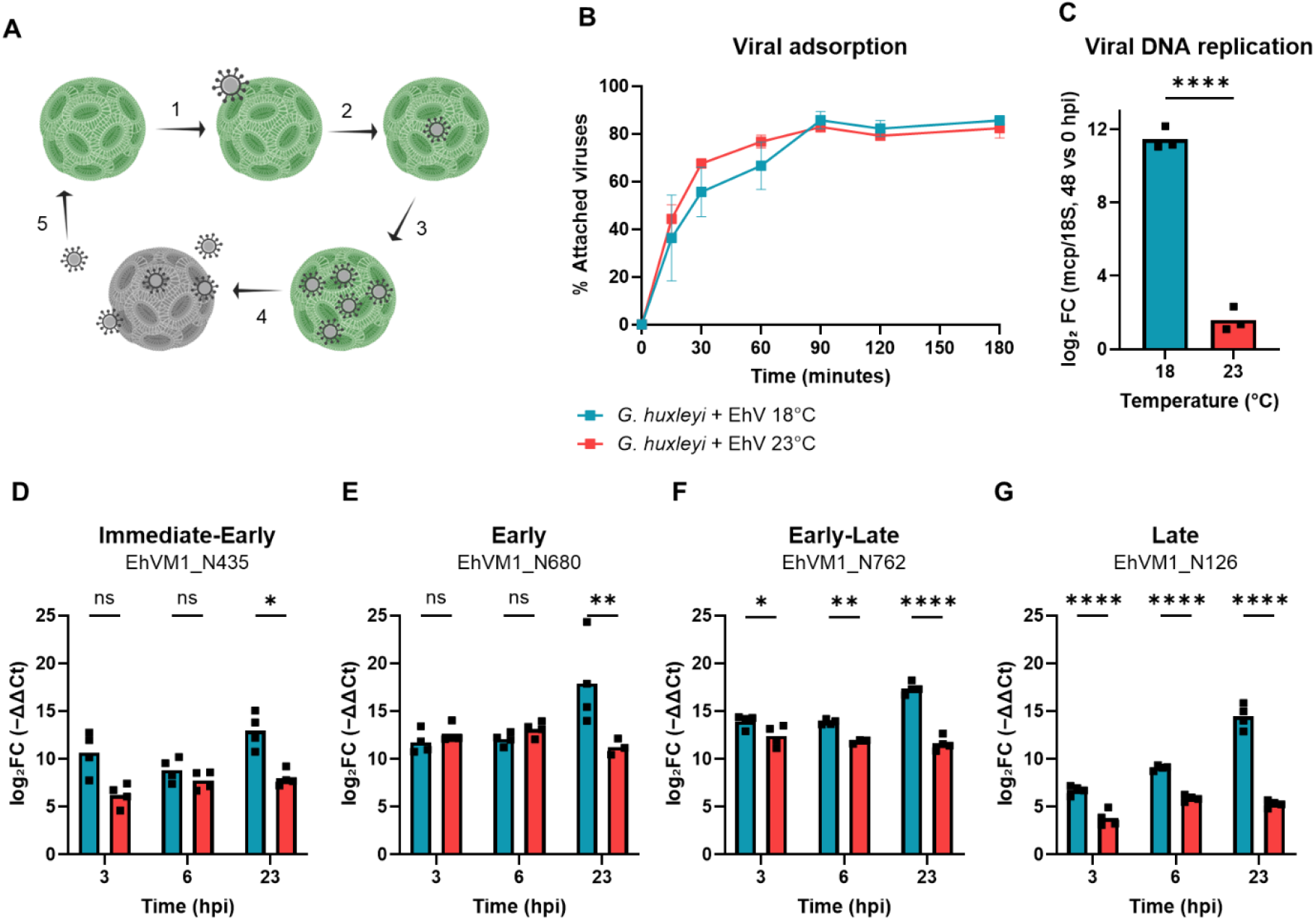
EhV infection is arrested at a distinct phase of its life cycle under elevated temperature. **A)** Schematic representation of the stages of viral infection. EhV adsorbs (1) and enters (2) a *G. huxleyi* cell. The cell becomes a virocell and supports viral transcription, translation, DNA replication and formation of progeny viral particles (3). After completion of the viral replication cycle, EhV virions are released into the surrounding water (4), where they can encounter other *G. huxleyi* cells, and infection can start anew (5). **B)** EhVM1 adsorption to *G. huxleyi* RCC6946 was assessed at 18°C (blue squares) and 23°C (red squares). The percentage of attached viruses was calculated as 100% minus the percentage remaining in the media (i.e., not adsorbed), as determined by flow cytometry and is presented as the average ± SD (n = 3, each averaged from three technical replicates). Error bars smaller than symbol size are not shown. **C)** Viral DNA replication was quantified in infected cultures at both temperatures. Replication is presented as log_2_FC (48hpi vs 0hpi) of the ratio of the viral major capsid protein (mcp) gene copies to the algal 18S gene copies, as quantified by digital droplet PCR. Statistical differences were tested using Welch’s t-test. **D-G)** Viral gene expression was assessed at 18°C (blue bars) and 23°C (red bars). Expression of representative viral genes from each kinetic class was quantified by RT-qPCR using the ΔΔCT method. The expression of each viral gene at each time point is normalized to the housekeeping gene comp74329_c0 and is relative to 0 hpi. Statistical differences (except for panel C) were tested using repeated measures two-way ANOVA followed by Sidak’s multiple comparisons test. p < 0.05 (*), p < 0.01 (**), p < 0.001 (***), p < 0.0001 (****). The scheme presented in (A) was created with Biorender.com. **Panels A-C can be used as the featured image**.

### Viral DNA replication is arrested at an elevated temperature

After confirming successful viral attachment at the elevated temperature, we sought to examine whether EhV can replicate its genome at 23°C. To assess DNA replication, we quantified the DNA copy numbers of the viral major capsid protein (mcp) gene and the algal 18S gene using digital droplet PCR. The ratio between mcp and 18S (hereafter mcp/18S) was used to normalize the viral gene copies to host gene copies. At 48 hpi at 18°C, the mcp/18S ratio increased by 11.5 log_2_-fold (corresponding to ∼2,800-fold increase) compared to 0 hpi (Fig. 3C). In contrast, at 48 hpi at 23°C, the mcp/18S ratio was significantly lower than at 18°C (p<0.0001), with only a 1.6 log_2_-fold (corresponding to ∼3-fold) increase compared to 0 hpi (Fig. 3C, Fig. S2A-B). Therefore, we conclude that the viral DNA replication is arrested at 23°C.

### Viral transcription is maintained at the early phase of infection but attenuated at subsequent stages under elevated temperature

We then continued to evaluate the viral replication cycle by assessing viral gene transcription. We followed the expression of representative genes from different kinetic classes by RT-qPCR. The genes belonging to the immediate-early and early kinetic classes showed similar transcription levels at both temperatures up to 6 hpi (Fig. 3D-E, S2C-D). By 23 hpi, log_2_FC of immediate-early genes was significantly lower at 23°C than at 18°C (8 vs 13 for EhVM1_N435 and 12 vs 14 for EhVM1_N485, p = 0.022 and p = 0.0206, respectively, Fig. 3D, S2C). The early viral gene EhVM1_N680 also had reduced expression at 23°C at 23hpi (log_2_FC of 11 vs 17, p=0.002, Fig. 3E). The early-late gene EhVM1_N758 log_2_FC was similar at both temperatures until 6 hpi but by 23 hpi was lower at 23°C (8 compared to 16, p<0.0001, Fig. S2E). All other early-late and late genes showed reduced expression as early as 3 hpi. The early-late gene EhVM1_N762 had log_2_FC of 12 vs 14 at 3 hpi, and 12 vs 18 at 23 hpi (23°C vs 18°C; p=0.0212, p<0.0001, Fig. 3F). The late gene EhVM1_N126 (mcp) had log_2_FC of 4 vs 7 at 3 hpi, and 5 vs 14 at 23 hpi (23°C vs 18°C; p<0.0001, p<0.0001, Fig. 3G). The late gene EhVM1_N170 had log_2_FC of 5 vs 8 at 3 hpi and 4 vs 7 at 23 hpi (23°C vs 18°C; p=0.0005, p<0.0001; Fig. S2F). Taken together, these results suggest that viral transcription is maintained at an elevated temperature during the early phase of viral infection but attenuated at subsequent stages.

### Viral infection arrest is reversible at the population level during a transient heatwave

We next aimed to test whether the halted infection under elevated temperature is permanent or can be reversed, in which a return to the basal conditions reactivates the infection. We infected *G. huxleyi* RCC6946 with EhVM1 at an MOP of 3 and placed the cultures at 18°C and 23°C. After 2 days, we moved the 23°C-infected cultures to 18°C. Within 4 days at 18°C, viral infection was restored as indicated by the 297-fold decline in algal abundance compared to the control cultures (p=0.0128, Fig. 4A), and the profound increase in viral abundance that reached similar levels to the 18°C-infected cultures (Fig. 4B). We then aimed to determine whether longer heat durations, resembling those of marine heatwaves, can alter infection outcomes. We infected *G. huxleyi* RCC6946 with EhVM1 and placed the samples at 23°C for 3 weeks. During the heatwave, algal abundance increased in both the control and infected samples. To maintain exponential algal growth, we diluted the cultures weekly with fresh media. In the first two dilutions, the infected cultures reached lower cell concentrations than the controls (1.6-fold at 5 dpi and 1.4-fold at 12 dpi, p=0.0029 and p=0.0054, respectively). At the third dilution (19 dpi), the infected culture reached a cell concentration similar to that of the control (Fig. 4C). Next, at 21 dpi, we transferred the cultures from 23°C to 18°C. Within 5 days (26 dpi), algal abundance declined by 187-fold compared to the control (p=0.0196, Fig. 4C). Accordingly, viral abundance increased by 2394-fold compared to the control (p=0.0427, Fig. 4D). Taken together, these results show that heat-induced infection arrest is reversible both for short- and long-term heat incubations at the population level.

**Figure 4.**
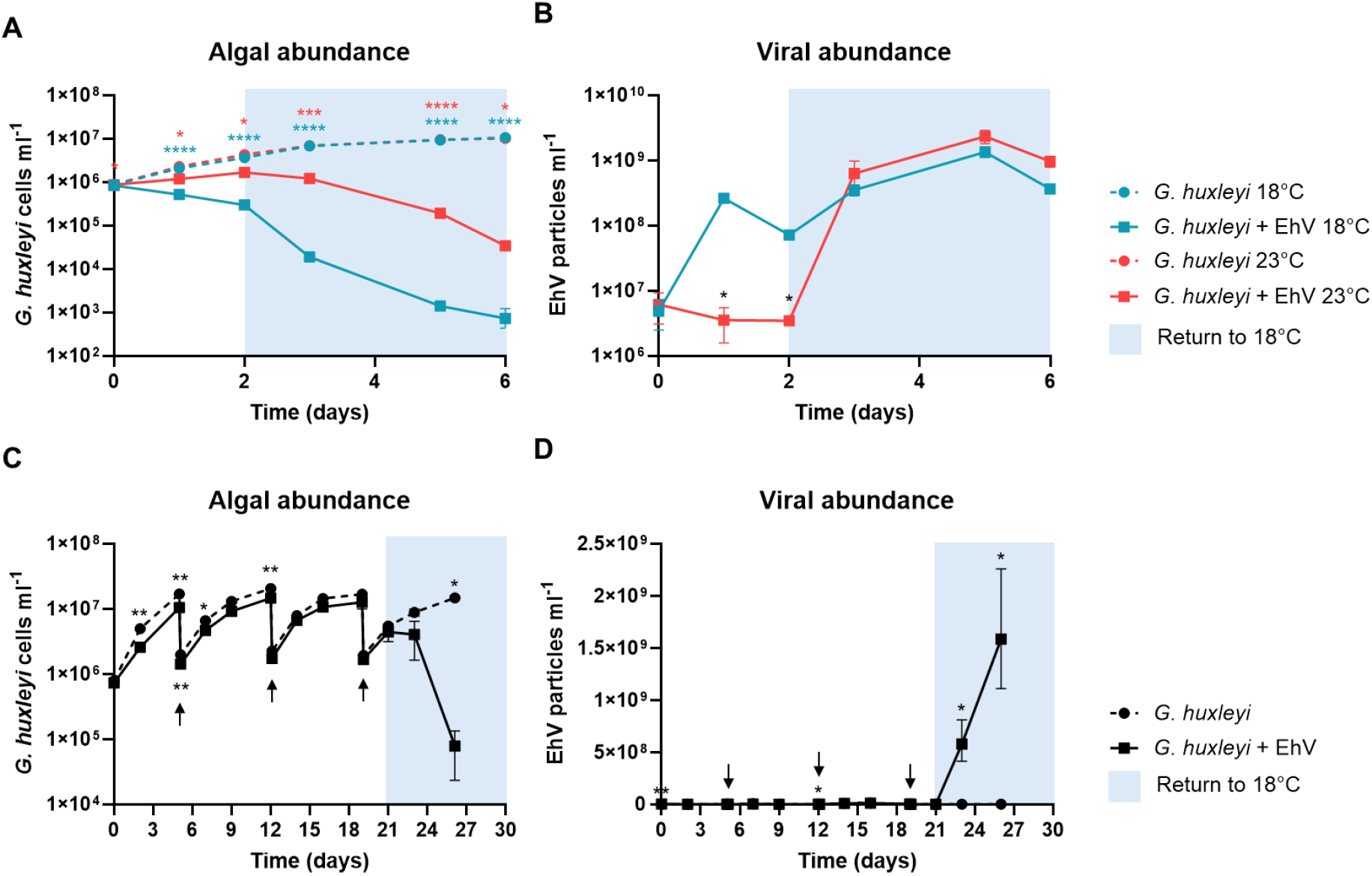
Heat-induced infection arrest is reversible at the population level. **A)** *G. huxleyi* RCC6946 was inoculated with EhVM1 at 18°C or 23°C. After 2 days, the heat-treated cultures (red squares) were moved to 18°C (depicted as a blue box). Algal abundance was evaluated for control and infected samples. **B)** Viral abundance during the experiment setup of A. **C)** Algal abundance in infected (solid line, square) and control (dashed line, circle) cultures during 3-weeks of incubation at 23°C. The cultures were diluted 1:10 into fresh media weekly (black arrows) to maintain exponential growth. After 21 days, the cultures were placed at 18°C (depicted as a blue box). **D)** Viral abundance during the experiment setup of C. Results were evaluated by flow cytometry and are presented as the average ± SD (n = 3). Error bars smaller than symbol size are not shown. Statistical differences were tested using a repeated-measures two-way ANOVA followed by Tukey’s multiple comparisons test between infected and control cultures with color representing temperature (A), 18°C-infected and 23°C-infected cultures (B) or Sidak’s multiple comparisons test between infected and control cultures (C, D). p < 0.05 (*), p < 0.01 (**), p < 0.001 (***), p < 0.0001 (****).

### Temperature-dependent infection arrest is reversible in a minor subpopulation of virocells

The reversibility of viral infection observed when the infected algal cultures were returned to 18°C after incubation at 23°C could be due to reactivation of infection within actively infected cells (virocells) or reinfection by extracellular virions remaining in the media. To distinguish between these two scenarios, we examined virocell infection dynamics at single-cell resolution. *G. huxleyi* RCC6946 was infected with EhVM1 and incubated at 18°C or 23°C for 22 hours. To assess whether individual infected cells can complete the infection cycle upon return to 18°C, we used a single-cell infectivity assay modified from Kirsner et al., 2016 [30]. Single cells from infected cultures were sorted into acceptor wells containing uninfected *G. huxleyi* RCC6946. The plates were then placed at 18°C. Lysis of cells in acceptor wells indicated that the sorted cells were virocells that completed their infection cycle upon return to 18°C (Fig. 5A). To increase the probability of sorting infected cells, we sorted single cells that displayed high autofluorescence in the green channel, as infected cultures displayed a shift toward higher green autofluorescence, possibly indicating stressed infected cells (Fig. S3A-B). Indeed, sorting high-green autofluorescence cells at 18°C increased the percentage of lysed acceptor wells by 1.6-fold (compared to sorting low-green autofluorescence cells, p=0.0025, Fig. S3C) and led to lysis in 69% of the wells within 6 days (Fig. 5B). In contrast, infected cells from 23°C led to the lysis of only 1.3% of the acceptor wells (p<0.0001, Fig. 5B). As a positive control for the presence of infected cells in the 18°C and 23°C cultures, we increased the probability of sorting virocells that can complete their infection cycle by sorting more than one cell into each acceptor well. Sorting 10 or 100 virocells from 23°C increased the lysis percentage in acceptor wells to 25% and 100%, respectively (Fig. S4). Accordingly, sorting 10 or 100 cells from 18°C increased the well lysis percentage to 100% (Fig. S4). To verify that lysis was due to infection by donor virocells rather than extracellular viruses remaining in the media, the virus-resistant strain *G. huxleyi* RCC379 was inoculated with EhVM1 at 18°C and underwent the same washing and sorting steps into exponentially growing *G. huxleyi* RCC6946 at 18°C. Notably, none of the wells were lysed (Fig. 5B), indicating that EhV was transferred to the acceptor wells only intracellularly, not via the culture media. To ascertain that the lysis in the acceptor wells was due to sorting of virocells and not other toxic effects, we sorted uninfected *G. huxleyi* RCC6946 from both 18°C and 23°C into acceptor wells, which yielded no lysis (Fig. 5B). Taken together, we conclude that only a small fraction of infected cells can resume infection when returned to basal temperature, suggesting that reinfection by extracellular viruses is the main driver of transiency of viral infection during a heatwave.

**Figure 5.**
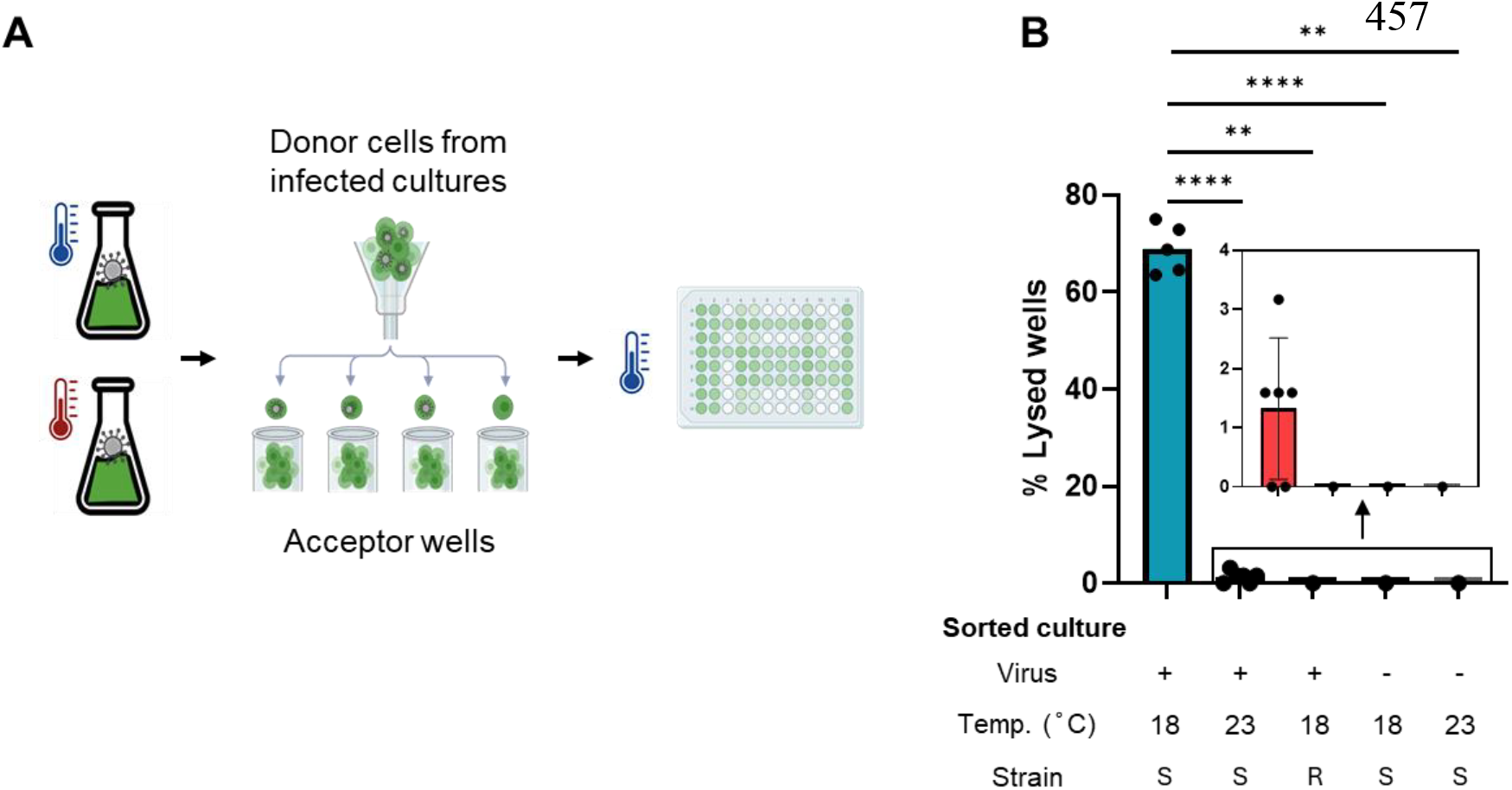
Temperature-dependent infection arrest is reversible in a minor virocell subpopulation. **A)** Schematic representation of the experiment to test reversibility from infection-arrest at the single-cell level. *G. huxleyi* RCC6946 and resistant *G. huxleyi* RCC379 cultures were inoculated with EhVM1 and placed at 18°C or 23°C for 22 hours. Single cells were sorted from the uninfected and infected cultures into individual wells of 96-well plates containing exponentially growing *G. huxleyi* RCC6946 at 18°C. Lysis of cells in acceptor wells indicated that the sorted cells were virocells that completed the infection cycle upon return to 18°C. Wells were considered lysed if their chlorophyll fluorescence, as measured by the plate reader, was < 500 (live wells had an average fluorescence signal > 5000). The resistant strain *G. huxleyi* RCC379 + EhVM1 was used as a control for the extracellular viruses in the media; 84 cells were sorted. For *G. huxleyi* RCC6946 *+* EhVM1 at 18°C or 23°C, 480 and 378 cells were sorted, respectively. For the *G. huxleyi* RCC6946 uninfected cultures, 96 cells were sorted from each temperature. **B)** The percentage of lysed wells in each plate 6 days post-sorting. The source of the sorted cells is depicted in the table (sorted culture). Infected culture (Virus +), control uninfected (Virus -), pre-sorting temperature (18°C vs 23°C), EhVM1 susceptible (Strain “S”) or resistant (Strain “R”) strains. The inset displays a magnified view of the boxed region. Statistical differences were tested using logistic regression. p < 0.01 (**), p < 0.0001 (****). The scheme presented in (A) was created with Biorender.com.

### Strain-specific response to viral infection under elevated temperature

To examine the heat-induced impairment of EhV infection of *G. huxleyi* in a broader ecological context, we tested EhVM1’s ability to infect several other *G. huxleyi* strains isolated from natural blooms. Interestingly, both *G. huxleyi* RCC6918 and *G. huxleyi* RCC6936, mesocosm isolates susceptible to EhVM1 at 18°C, did not lyse at 23°C (Fig.6A-B). We further tested the effect of elevated temperature on viral infection using different host-virus pairs. *G. huxleyi* CCMP2090 or *G. huxleyi* CCMP374 were inoculated with EhV201 at 18°C and 23°C. At both temperatures, the virus lysed *G. huxleyi* CCMP2090, although algal demise was slower at 23°C, with the final density 4.5-fold higher than at 18°C (p=0.0018, Fig. 6C). Interestingly, EhV201 did not completely lyse *G. huxleyi* CCMP374. The final algal density was 62-fold higher than that of 18°C-infected culture (Fig. 6D). Taken together, these results indicate differences in the temperature sensitivity of viral infection across different host-virus pairs.

**Figure 6.**
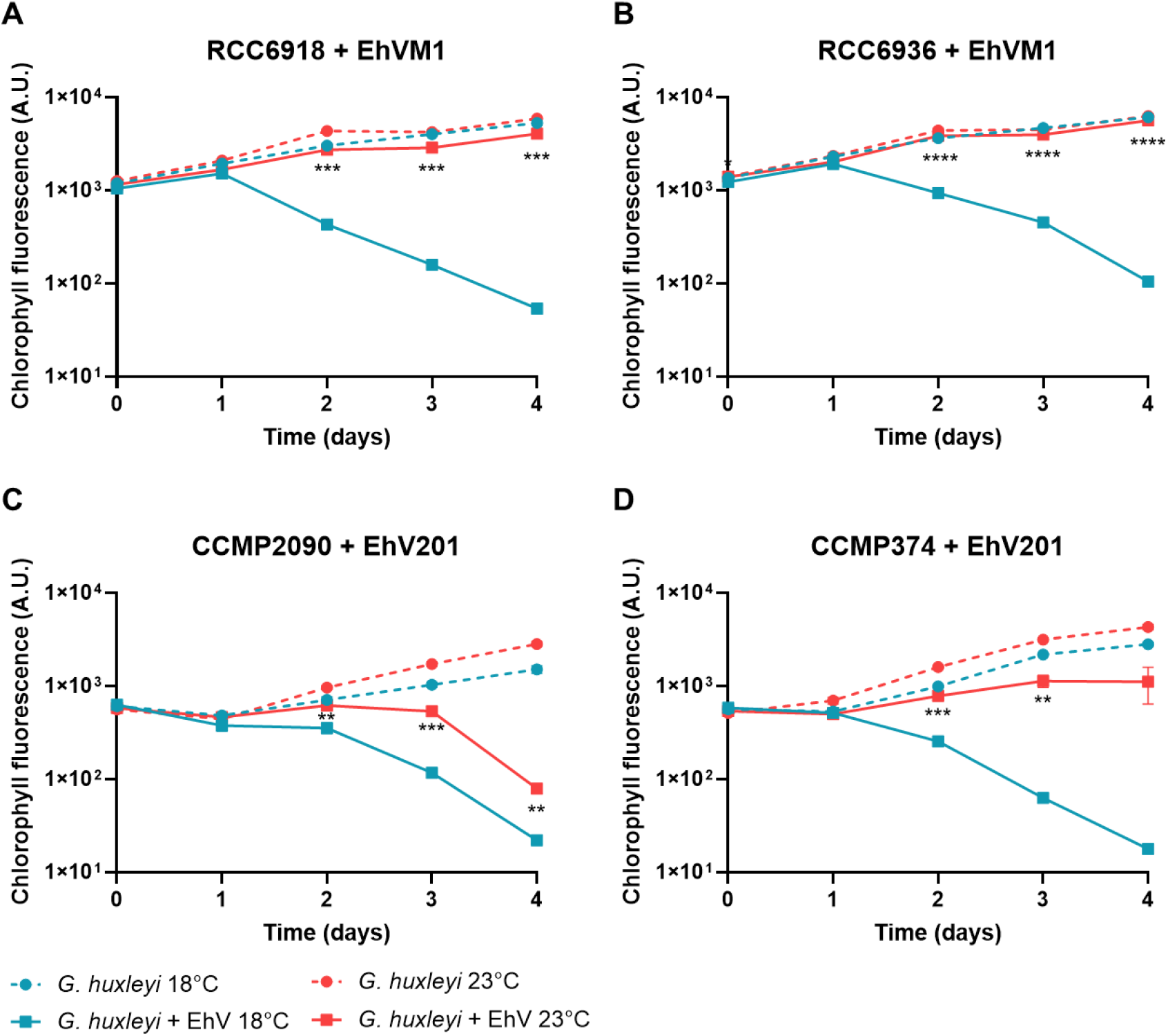
Sensitivity of viral infection to elevated temperature varies among host-virus pairs. Pairs of *G. huxleyi* and EhV strains were inoculated at 18°C (blue squares) and 23°C (red squares). EhVM1 was inoculated with *G. huxleyi* RCC6918 **(A)** or *G. huxleyi* RCC6936 **(B)**. EhV201 was inoculated with *G. huxleyi* CCMP2090 **(C)** or *G. huxleyi* CCMP374 **(D)**. The infection dynamics were monitored by measuring chlorophyll fluorescence in the cultures using a plate reader. Results represent average ± SD (n = 4). Error bars smaller than symbol size are not shown. Statistical differences were tested using a two-way repeated-measures ANOVA followed by Tukey’s multiple comparisons test of 18°C infected vs 23°C infected. p < 0.05 (*), p < 0.01 (**), p < 0.001 (***), p < 0.0001 (****).

## Discussion

Anthropogenic greenhouse gas emissions have led to increased heat and carbon dioxide uptake by the ocean, causing accelerated ocean warming and acidification [31-34]. Rising temperatures increase upper ocean stratification, suppressing vertical mixing, thus impacting the vertical exchanges of heat, carbon, dissolved oxygen, and nutrient availability [35, 36]. These environmental shifts are expected to reshape microbial community diversity and distribution [37-39]. Despite extensive research on individual and combined stressors affecting species and community composition, the consequences of climate change for microbial interactions, including host–virus systems, remain largely unknown. In this work, we have described a virocell-specific sensitivity to an elevated temperature. Upon heat, the virus enters the algal host cell, followed by induction of early viral gene expression, but it cannot complete its transcriptional program or replicate its genome (Fig. 3). Following heat exposure, a return to basal temperature enables infection to resume at the population level, even after prolonged exposure of 21 days (Fig. 4). At the single-cell level, only a minor subpopulation of infected cells could complete the infection that was halted by the heat treatment (Fig. 5). We further showed that the sensitivity of viral infection to elevated temperature varied among host-virus pairs (Fig. 6).

Impaired infection can result from loss of virion infectivity, induction of host resistance mechanisms, or failure of a process within the virocell that is crucial for the viral life cycle. We demonstrated that pre-exposure of virions to 23°C for at least 72 hours did not result in loss of viral infectivity (Fig. 2), consistent with previous studies which showed that EhV86 infectivity is not compromised by exposure to an elevated temperature of 21°C [24] or even a brief exposure to an extreme temperature of 40°C [40]. This demonstrates the EhV virion’s ability to survive changes in seawater temperature and remain infective under varying environmental conditions. Alternatively, infection arrest can be caused by induction of host resistance to viral infection, which can be achieved by blocking infection at different stages [41] and activating defense strategies [29]. Here, we showed that pre-exposure of the algal cells to elevated temperature does not induce host resistance (Fig. 2), suggesting that elevated temperature alone does not induce a host immune response prior to infection. Nevertheless, we cannot exclude a possible explanation in which resistance mechanism is induced only upon infection. Resistance can also occur when the host is compromised in a molecular component essential for viral infection, such as a receptor, thereby conferring extracellular resistance and eliminating the virus’s ability to adhere to cells. In contrast to a previous study [24], we showed that the halted infection during heat exposure is not due to extracellular resistance. EhVM1 successfully attaches to and enters *G. huxleyi* RCC6946, but once within the cell, it cannot complete its replication cycle, suggesting that a specific phase of the infection cycle is compromised at the subcellular level (Fig. 3). A similar form of intracellular resistance, although not caused by heat stress, was described in the *Chlorella variabilis* NC64A*-*chlorovirus OSy-NE5 system. During infection, OSy-NE5 entered the cells and initiated infection but could not replicate its DNA, leading to the release of some empty, noninfectious viral particles [42]. Viral infection is not a synchronous process, as individual cells can differ in their infection state [13]. However, heat treatment completely halted infection, suggesting that an essential enzyme that acts at the transition to the mid-to late stages of infection may be heat-sensitive. Notably, EhV encodes its own DNA and RNA polymerases, possibly enabling genome replication and transcription independent of the host machinery [13]. We showed that viral DNA replication is blocked at 23°C (Fig. 3), possibly suggesting that the viral DNA polymerase is heat-sensitive. Interestingly, there are known examples of temperature sensitivity of the DNA polymerase of other giant viruses, such as *adenovirus* and *poxvirus*, with mutants exhibiting defective DNA synthesis at non-permissive temperatures [43, 44]. Furthermore, in many dsDNA viruses, active DNA replication is required for late gene transcription [45-47], supporting our findings of attenuation of the late infection stage.

Marine heatwaves can persist for several days to several months, with peak sea surface temperature anomalies of up to 5°C [4, 16]. *G. huxleyi* blooms can persist for two to three months in high latitudes [48]. We demonstrated in the lab that viral infection was arrested during an ecologically relevant timeframe of a three-week heat treatment (Fig. 4C, D). This infection arrest may have important ecological and biogeochemical implications in the ocean, as algal mortality by viral infection diverts carbon flux from higher trophic levels to bacterial respiration and deep-ocean sequestration [7, 8, 49]. Furthermore, arrest of viral infection in a natural *G. huxleyi* bloom may alter bloom duration and enhance the impact of other mortality agents in controlling bloom termination. Apart from viruses, bacteria, grazers, and environmental factors collectively regulate algal bloom dynamics and their demise [7, 50-55]. Ocean warming can decrease zooplankton grazing rates on bacteria and phytoplankton [56]. Additionally, warming may induce bacterial pathogenicity toward phytoplankton [57, 58]. We have shown that following heat exposure, a return to basal temperature enables viral infection to resume at the population level. This reversibility is likely due to reinfection of the algal culture by residual extracellular viruses that remain infective after heat exposure (Fig. 2, 4). Thus, in the context of natural blooms, the timing of a marine heatwave might dictate whether infection persists. A marine heatwave that occurs at the start of a bloom may not abolish infection that occurs at the late bloom stage. Interestingly, at the single-cell level, 1.3% of cells exposed to heat were able to complete the infection upon return to 18°C (Fig. 5B). In the ocean, this small percentage of cells that already harbor viruses and can recover from infection arrest may be better fitted to act as a seed for viral propagation than virions that need to successfully encounter new cells. Additionally, this raises the question of the fate of the infected cells at 23°C that could not continue the infection when returned to 18°C, and whether they are ultimately cured of viral infection.

Viral infection sensitivity to temperature was strain-specific and varied among host-virus pairs. All *G. huxleyi* strains that were susceptible to EhVM1 infection at 18°C showed an inhibition of infection at 23°C (Fig 1, Fig. 6A-B). In contrast, the effect of elevated temperature on EhV201 infection varied among algal hosts. EhV201 lysed *G. huxleyi* CCMP2090 at both temperatures, whereas it did not lyse *G. huxleyi* CCMP374 at 23°C (Fig.6 C-D). As *G. huxleyi* exhibits wide intraspecific thermal diversity, with the thermal niche width ranging from 6°C to 25°C [15, 16], it stands to reason that different strains of its specific virus also vary in their thermal response mechanisms. Similarly, the distribution of *Micromonas* virus (MicV) in the surface ocean is largely temperature-driven, with different viral strains harboring various thermal optima [59]. Moreover, the lytic cycle of MicV is linked to optimal *Micromonas* growth at 20 to 25 °C. Above these optimal temperatures, lytic infection is significantly reduced [23]. The combined response of *G. huxleyi* and EhV to temperature will determine whether the infection is successful. Furthermore, viral pressure may shape host thermal response curves, especially if adaptation to higher temperatures confers resistance to viral infection. Nevertheless, according to the evolutionary arms race theory [60], it is possible that future EhVs will overcome heat sensitivity, or that viruses less affected by elevated temperatures will become more prevalent.

Understanding the temperature-driven changes in host-virus interactions is critical for predicting the future of oceanic carbon flux and ecosystem stability. By investigating the sensitivity of distinct phases of viral infection to elevated temperature, we discovered an intracellular form of infection arrest. A valuable approach to understanding the molecular mechanism underlying this phenomenon is to examine the heat sensitivity of the viral metabolic processes and enzymatic activities. We propose that the reversibility of infection arrest is a key mechanism for the recovery of *G. huxleyi*-EhV interactions from marine heatwaves.

## Acknowledgements

We are grateful to Dr. Shifra Ben-Dor and Dr. Ron Rotkopf for their assistance with data analysis. We thank Ian Probert and Martin Gachenot from the Roscoff Culture Collection for isolating the *G. huxleyi* strains during the mesocosm experiment. This research was supported by the European Research Council AdV (VIBES, grant no. 101053543) and by the Institute for Environmental Sustainability (IES) at the Weizmann Institute of Science, both awarded to A.V.

## Author Contributions

Noa Shima, Daniella Schatz, and Assaf Vardi (conceptualizing the study and experimental design), Noa Shima (performing the experiments), Talia S. Shaler (assigning predicted kinetic classes to EhVM1 genes), Daniella Schatz (technical assistance in the ddPCR experiment), Nir Joffe (technical and conceptual assistance in the single-cell sorting experiment), Noa Shima and Assaf Vardi (writing the manuscript), Nir Joffe, Talia S. Shaler, and Daniella Schatz (reviewing and editing the manuscript).

## Conflicts of interest

None declared.

## Data availability

All data generated or analysed during this study are included in this published article [and its supplementary information files].

## Notes

### Competing Interest Statement

The authors have declared no competing interest.

### Summary of Updates

The author list was updated to correct order, and row numbers were added.

