## Supplementary figures 1-4 for "Reversible inhibition of viral life cycle in response to elevated temperature in a bloom-forming alga"

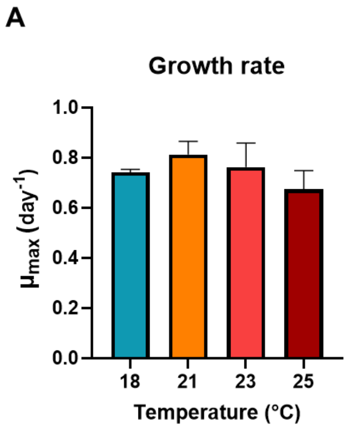


**Figure S1. Effect of temperature on *G. huxleyi* cell growth.**
**A)** Algal growth rate ($\mu_{max}$) of uninfected cultures at 18˚C, 21˚C, 23˚C, and 25˚C. Results represent average ± SD (n = 3). Statistical differences were tested using one-way ANOVA followed by Tukey's multiple comparisons test.

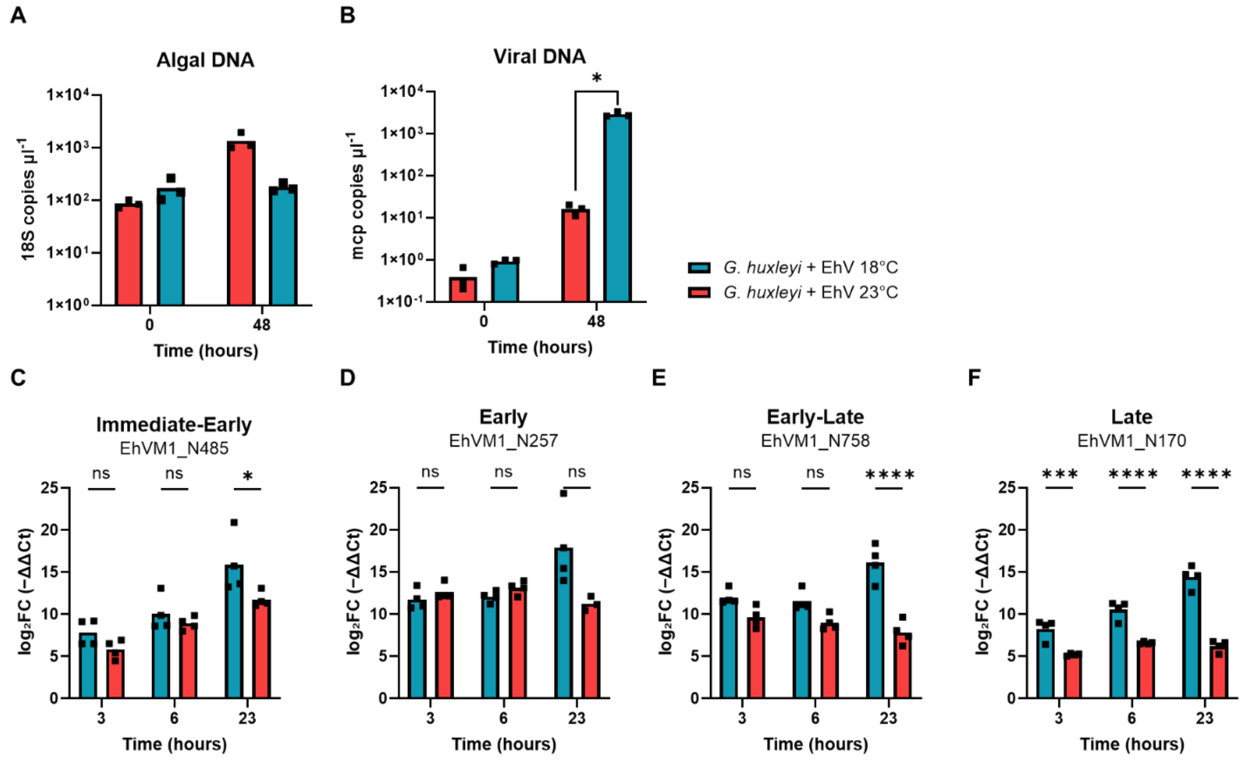


**Figure S2. Viral DNA replication and late gene transcription are arrested under heat stress.**
**A)** Algal 18S DNA copies in infected cultures at 18˚C (blue bars) and 23˚C (red bars). The number of 18S gene copies was evaluated using digital droplet PCR. **B)** Viral mcp DNA copies as evaluated by digital droplet PCR. **C-F)** Viral gene expression at 18˚C (blue bars) and 23˚C (red bars) as evaluated by RT-qPCR. Expression of representative viral genes from each kinetic class was quantified using the ΔΔCT method. The expression of each viral gene at each time point is normalized to the housekeeping gene comp74329_c0 and is relative to 0 hpi. Results represent

average ± SD (n = 3). Error bars smaller than symbol size are not shown. Statistical differences were tested using a repeated-measures two-way ANOVA followed by Sidak's multiple comparisons test. $p < 0.05$ (*), $p < 0.001$ (***), $p < 0.0001$ (****).

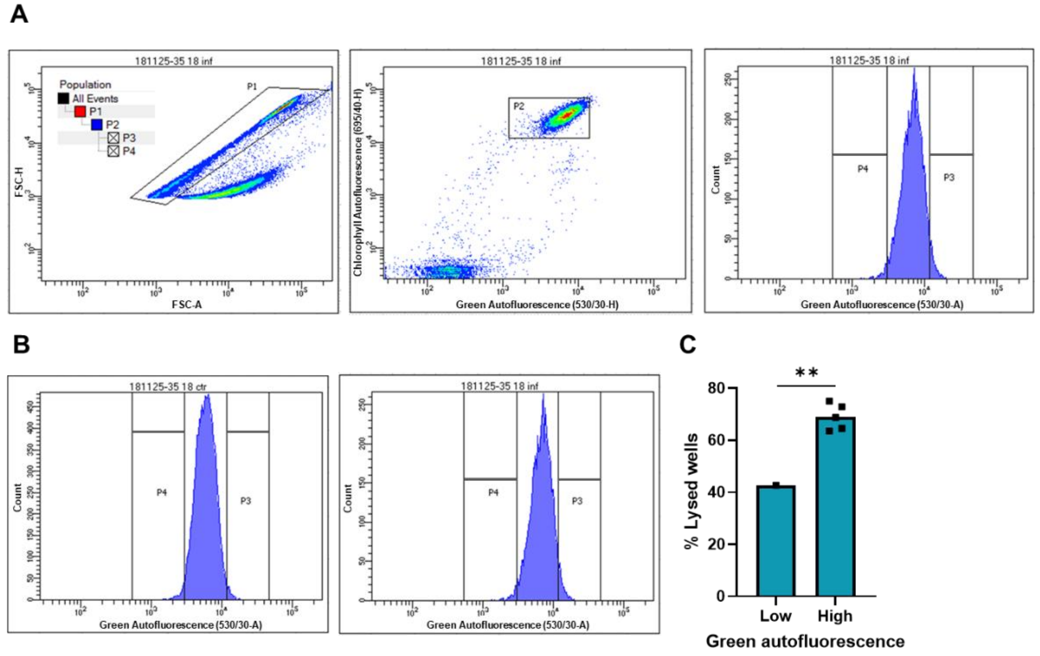


**Figure S3. Sorting cells with high green autofluorescence increases well lysis percentage. A)** Representative plots showing the gating strategies for choosing single cells for sorting. Single cells (P1 gate) were chosen by their FSC-A (x-axis) and FSC-H (y-axis) ratio (left dot plot). *G. huxleyi* main population (P2 gate) was chosen from high chlorophyll events and autofluorescence in the green channel (x-axis 488-530/30-A, y-axis 488-695/40-H, middle dot plot). Then, cells with high (P3 gate) or low (P4 gate) autofluorescence in the green channel (x-axis 488-530/30-A, histogram on the right) were selected for sorting. Fluorescence is measured in arbitrary units. **B)** Histogram of autofluorescence in the green channel (x-axis 488-530/30-A) of uninfected (left) and infected (right) *G. huxleyi* RCC6946 18˚C culture. C) Percentage of well lysis in each plate 6 days post-sorting low (96 wells) or high (480 wells) green autofluorescent cells from *G. huxleyi* RCC6946 18˚C-infected culture. Statistical differences were tested using logistic regression. $p < 0.01$ (**).

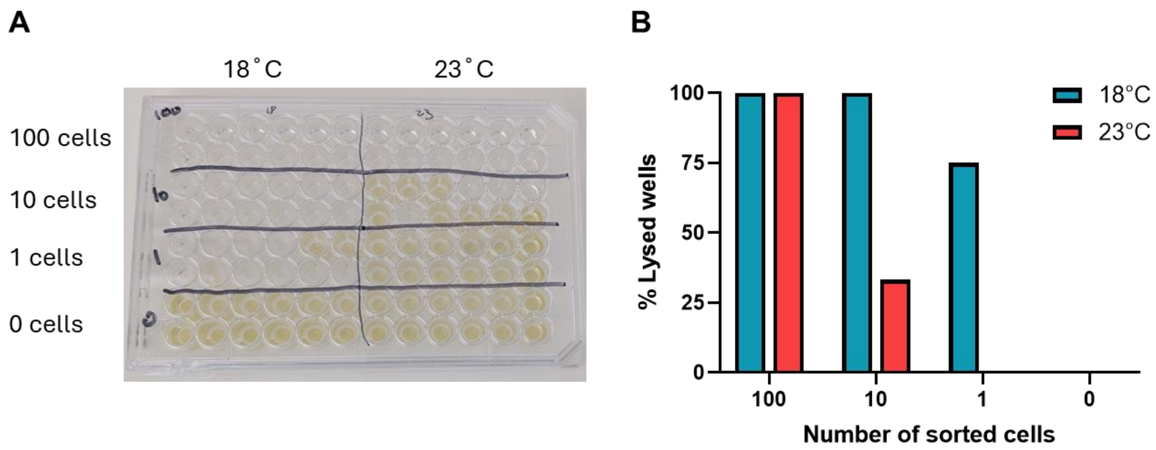


**Figure S4. Sorting higher cell numbers into each well increases well lysis percentage. A)** One representative 96-well plate, 6 days post-sorting of different numbers of single cells from infected cultures, is shown. Cells from 18˚C were sorted to the left half of the plate. Cells from 23˚C were sorted to the right half of the plate. No cells were sorted into the 2 bottom rows, as a control for the algal growth in the plate. The rows above contained wells with 1, 10 or 100 sorted cells (bottom up). **B)** Percentage of wells lysed when inoculated with different numbers of single cells, as shown in A. n=12 wells per treatment.
